# A machine-learning classifier trained with microRNA ratios to distinguish melanomas from nevi

**DOI:** 10.1101/507400

**Authors:** Rodrigo Torres, Ursula E Lang, Miroslav Hejna, Samuel J Shelton, Nancy M Joseph, A. Hunter Shain, Iwei Yeh, Maria L. Wei, Michael C Oldham, Boris C Bastian, Robert L Judson-Torres

## Abstract

The use of microRNAs as biomarkers has been proposed for many diseases including the diagnosis of melanoma. Although hundreds of microRNAs have been identified as differentially expressed in melanomas as compared to benign melanocytic lesions, limited consensus has been achieved across studies, constraining the effective use of these potentially useful markers. In this study we quantified microRNAs by next-generation sequencing from melanomas and their adjacent benign precursor nevi. We applied a machine learning-based pipeline to identify a microRNA signature that separated melanomas from nevi and was unaffected by confounding variables, such as patient age and tumor cell content. By employing the ratios of microRNAs that were either enriched or depleted in melanoma compared to nevi as a normalization strategy, the classifier performed similarly across multiple published microRNA datasets, obtained by microarray, small RNA sequencing, or RT-qPCR. Validation on separate cohorts of melanomas and nevi correctly classified lesions with 83% sensitivity and 71-83% specificity, independent of variation in tumor cell content of the sample or patient age.

## Introduction

Misdiagnosis of cutaneous melanoma is among the most significant contributors to medical malpractice lawsuits in the United States (Wallace et al. 2013). The advanced stages of melanoma are associated with five-year survival rates less than 20% and have been responsible for over 10,000 deaths in the U.S. each year (Gershenwald et al. 2017; https://www.cancer.org/cancer/melanoma-skin-cancer/about/key-statistics.html). Although the disease is curable when detected and treated early, the process of differentiating between malignant lesions and the more prevalent benign lesions is challenging. The definitive diagnosis of concerning lesions is achieved through histopathologic assessment of a biopsy specimen, but a considerable rate of discordance even among expert pathologists has been established (Heenan et al. 1984; Boiko et al. 1994; Corona et al. 1996; Farmer et al. 1996; Brochez et al. 2002; Shoo et al. 2010; Gaudi et al. 2013; Niebling et al. 2014; Elmore et al. 2017; Elder et al. 2018). Although accuracy improved following implementation of more defined diagnostic criteria, a large-scale study published by Elmore and colleagues in 2017 reported interobserver discordance rates as high as 57-75% and intraobserver discordance rates at 37-65% (Elmore et al. 2017). Together, these observations highlight the complexity and subjectivity of histopathologic assessment and emphasize the need for objective methods for distinguishing malignant from benign lesions to augment current practices.

Molecular biomarkers can provide robust, objective and quantitative measurements of disease state (Rodríguez-Cerdeira et al.; Leachman et al. 2017; Buchbinder and Flaherty 2016). One class of candidate biomarkers is small non-coding microRNAs (miRNAs). Discovered twenty-five years ago (Lee et al. 1993), miRNAs stabilize transcriptional programs (Ebert and Sharp 2012; Judson et al. 2013) and their expression can distinguish cell state transitions during mammalian development and in disease progression (Parchem et al. 2014; Reddy 2015). Combined with a smaller size, reduced complexity, and superior stability over mRNA transcripts, miRNAs are appreciated as potentially valuable candidate biomarkers for many conditions and diseases (Jung et al. 2010; Sheinerman and Umansky 2013). However, despite abundant studies focused on a breadth of diseases, few miRNA biomarkers have emerged in the clinical setting (Pogribny 2018). One reason these promising candidate biomarkers have yet to reach their potential is the frequent lack of reproducibility between differential expression studies (Mumford et al. 2018; Nair et al. 2012; Raya et al. 2012; Witwer and Halushka 2016). Discrepancies in differential expression signatures across comparable studies have been attributed to sample heterogeneity, platform-specific biases in miRNA detection, and an absence of standardized normalization strategies (Mumford et al. 2018; Raya et al. 2012; Witwer and Halushka 2016).

Exemplifying these complications are studies that have explored the use of miRNAs as biomarkers for melanoma (reviewed in (Jarry et al. 2014; Jayawardana et al. 2016; Margue et al. 2013; Raya et al. 2012)). Independent studies using various platforms (microarray, RT-qPCR array, RNA-seq) to compare miRNA profiles between benign melanocytic lesions and melanomas have resulted in more than 500 different miRNAs identified as significantly differentially expressed (summarized in Tables S1 & S2) (Xu et al. 2012; Jukic et al. 2010; Sand et al. 2013; Komina et al. 2016; Kozubek et al. 2013; Hanniford et al. 2015; Latchana et al. 2017; Chen et al. 2011). However, only seven of these miRNAs showed reproducible expression differences in at least half of the cohorts, and none were identified in every study (summarized in Figure S1). Several of the most reproducibly identified miRNAs – miR-211-5p, miR-125b-5p, and miR-21-5p – have been validated as differentially expressed in benign and malignant pigmented lesions using in situ hybridization, suggesting some miRNAs could function as biomarkers (Babapoor et al. 2016; Wandler et al. 2017). However, differential expression of these same miRNAs was not observed in 10-30% of cohorts. Further investigations into the causes of these inconsistencies, and potential solutions, are needed.

In this study, we sought to determine whether a miRNA signature can reliably distinguish malignant from benign melanocytic lesions across both published and independently generated datasets. Machine learning-based classification can help distinguish predictive features from confounding variables, provided the model is trained on an appropriately controlled and annotated dataset (Guyon and Elisseeff 2003). We generated such a dataset and employed methods of machine learning to both identify the most common confounding variables influencing quantification of miRNA expression from FFPE samples and to generate a refined miRNA signature that correlated uniquely with diagnosis.

## Results

### Variation in tumor cell content of FFPE samples confounds miRNA expression analyses

To identify covariates that could confound differential miRNA expression analyses, we took advantage of a cohort of primary melanomas with intact adjacent benign nevi, from which they arose (Shain et al. 2018) (Fig. 1a). The different progression stages for each sample were diagnostically classified by a panel of at least five dermatopathologists and micro-dissected and genotyped for over five hundred cancer-related genes. Phylogenic trees of the somatic mutations identified in the respective tumor areas were constructed and confirmed the common clonal origin for the different progression stages of each patient. We estimated the tumor cell content (referred to here as tumor cellularity) using allele frequencies and magnitudes of copy number changes as previously described (Shain et al. 2018). Consequently, the dataset was annotated with both clinical features (e.g. patient age, sex, anatomical location of the lesion) as well as genomic information (e.g. mutation burden, copy number variation, tumor cellularity) for each matched pair of nevus and melanoma regions (Fig. 1b and Table S3).

**Figure 1:**
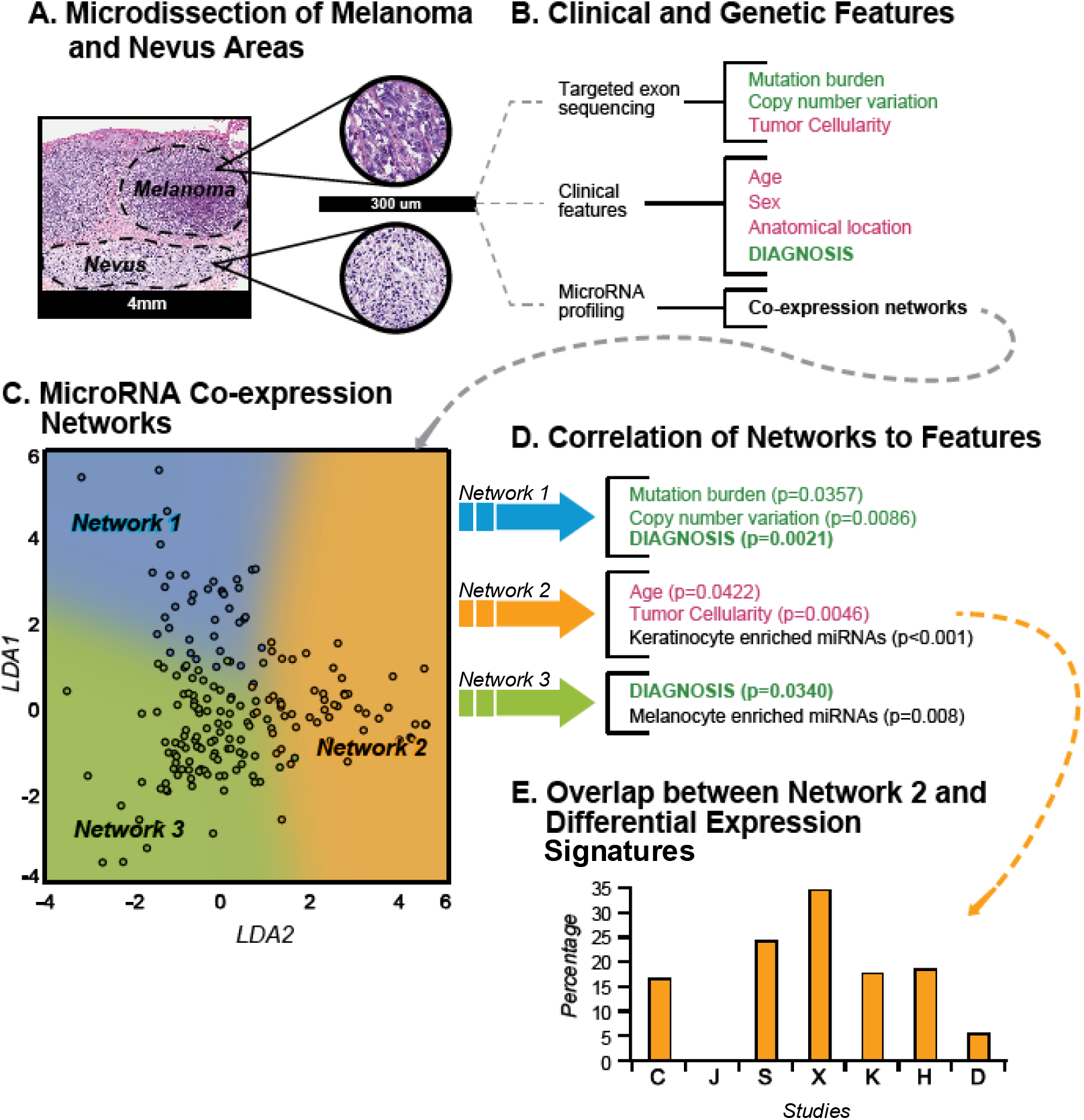
Tumor cellularity and age confound miRNA profiling of melanoma samples A) A cohort of melanomas with intact adjacent precursor nevus were identified and micro-dissected. B) List of features obtained for each micro-dissected region either from targeted exon sequencing (genetic features) or the pathology requisition form (clinical features). Red font indicates potentially confounding features. Green font indicates target feature (diagnosis) or genetic features highly correlated with target feature. C) miRNA co-expression networks. Scatter plot of all expressed miRNAs separated by LDA trained on the three networks. The three coexpression networks are indicated by the color of the points (blue, orange, and green). D) Correlations of PC1 of each miRNA network expression matrix with clinical and genetic features from (B) with p-values calculated from the corresponding correlation coefficient. E) Percent of miRNAs identified as differentially expressed between nevi and melanoma in seven published studies that overlap with Network 2. Published studies are identified by the first letter of the first authors’ last names (Table S1).

To investigate the influence of each genomic and clinical feature on the miRNA expression pattern, we conducted miRNA sequencing on fifteen of the regions from seven cases (Table S3). In order to first identify potential systemic confounding features, we first employed co-expression analyses for identification of networks of miRNAs sharing similar expression patterns across all regions and identified three co-expression networks (Fig. S2a, Table S4) that were effectively separated via Linear Discriminate Analysis (LDA) (Fig. 1c) (Langfelder and Horvath 2008). Each network consists of miRNAs with read counts that are positively correlated across all samples, regardless of level of expression. We next sought to determine whether the expression patterns of these networks correlated with any of the clinical or genomic annotations of the samples, including not only diagnosis but also potentially confounding features, such as patient age. We summarized the miRNA expression matrix for each network by its first principal component and compared these to the sample covariates (Fig. 1d, Fig. S2b-c). Two of the networks (Network 1 and Network 3) were significantly correlated with a diagnosis of melanoma and were not influenced by tumor cellularity or any other clinical feature. Network 1 was also correlated with mutation burden and copy number variation, both measurements of genome damage that increase during progression from nevus to melanoma (Shain et al. 2015).

In contrast to the two melanoma-associated networks, Network 2 was positively correlated with tumor cellularity and, to a lesser extent, patient age. This observation suggests that although miRNAs within Network 2 were differentially expressed in melanoma and nevus samples, variation in their observed abundance may reflect the extent of contamination with non-tumor cells rather than different progression stages. Consistent with this interpretation, we observed that miRNAs known to be expressed in cultured primary human keratinocytes were enriched in Network 2 as would be expected if keratinocytes were a significant fraction of contaminating non-tumor cells (Fig. 1d, Fig. S2c). Conversely, miRNAs known to be expressed in cultured primary human melanocytes were enriched in Network 3 consistent with changes in Network 3 reflecting melanocyte biology. Together, these data suggest that miRNA profiling datasets derived from micro-dissected FFPE samples can contain sufficient levels of contaminating non-tumor cells to influence the overall miRNA expression profile. Contamination by non-tumor cells is expected to vary among samples dependent on their size, histologic type (predominantly junctional versus intradermal), and preparation (e.g. precision of microdissection). If not controlled for, variation in tumor cellularity is expected to degrade the reproducibility of signatures across studies. Indeed, miRNAs from Network 2 constituted up to thirty percent of the miRNAs in expression signatures reported from the seven previously reported datasets (Fig. 1e). This result highlights the need for alternative analytical methods for identifying miRNA signatures predictive of melanoma diagnosis from FFPE derived samples.

### Classification of nevus from melanoma samples with miRNA ratios

To identify miRNAs that best distinguish malignant from benign melanocytic lesions, we utilized feature selection (FS). FS is a method of machine learning that is frequently used to simplify predictive models and to avoid analytical pitfalls such as the phenomena of over-fitting and the ‘curse of dimensionality’ (He and Yu 2010; Saeys et al. 2007). In the context of biology, FS methods can be applied to gene expression datasets to identify sets of features (in this case, miRNAs) that are more often biologically relevant and ultimately improve classification performance (Abeel et al. 2010). There are common FS methods, such as univariate statistical test filtering (e.g. FDR, t-test) and feature rank wrappers (e.g. backward selection) that will identify individual features that are independently relevant, but they miss features that are only relevant in the context of complex networks (Mnich and Rudnicki 2017; Wenric and Shemirani 2018). An alternative strategy is the all-relevant features (ARF) selection approach that involves multiple iterations of feature ranking and can determine both weak and strong relevant features (Kursa and Rudnicki 2010). An ARF method called Boruta (Fig. S3) has been shown to provide improved performance in a variety of datasets including gene array data and environmental science data (Li et al. 2016; Kursa 2014).

We employed Boruta to obtain an initial list of those miRNAs that were most important for discriminating the benign and malignant regions from our cohort across 1000 random forest iterations (Fig. 2a). All miRNAs with more than five total reads were considered, resulting in 341 unique features. For each miRNA, a second artificial feature was generated through randomized re-distribution of the read counts across samples (Fig. S3). These ‘shadow features’ provided an equal number of negative control features for which to compare each experimental feature. We conducted Boruta with the combined 682 experimental and negative control features, ranking the importance of each feature for the accurate classification of nevus samples from melanoma samples with each iteration. We identified 38 miRNAs that ranked higher than the maximum-performing shadow feature with a p-value of less than 0.001 (Fig S4). To enable comparison with published studies we also required that the expression levels of each miRNA were assessed in all published datasets (Table S2). The final list of feature-selected miRNAs contained two miRNAs with increased expression (miR-31-5p, miR-21-5p) and four miRNAs (miR-211-5p, miR-125a-5p, miR-125b-5p, miR-100-5p) with decreased expression in melanomas (Fig. 2c). These miRNAs are referred to as melanoma-enriched and melanoma-depleted miRNAs, respectively.

**Figure 2:**
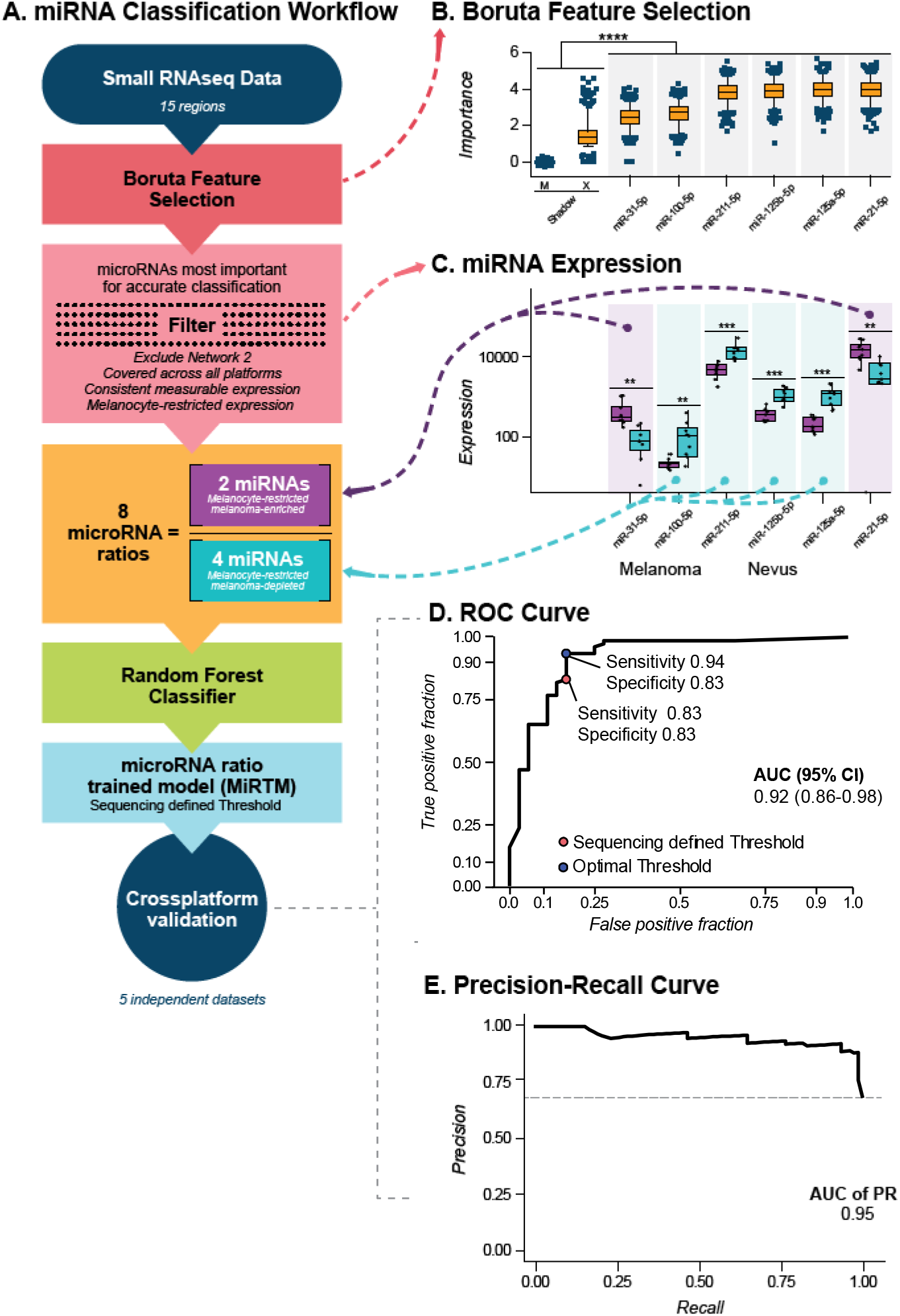
The development of a miRNA ratio-trained model that classifies melanocytic lesions A) Workflow of miRNA-Seq classifier development and testing. B)Top miRNAs for classifying nevi from melanomas using Boruta feature selection. Each feature-selected miRNA has a significantly higher importance value than the random shadow max feature (X) and shadow mean feature (M) (Fig. S3). The top six miRNAs are shown (full list in Fig. S4). C) Normalized miRNA-Seq counts from micro-dissected FFPE samples. Counts from melanoma regions (purple) and nevus regions (blue) are shown. Boxes indicate mean, first and third quartiles. miRNAs designated as melanoma-enriched or melanoma-depleted are designated by light purple and light blue backgrounds, respectively. D) ROC curves for cross-platform testing of MiRTM using the combined set of publicly available datasets. Threshold determined from the discovery sequencing set and optimal threshold are shown in as red and blue points respectively with corresponding sensitivity and specificity annotated for each. E) Precision-Recall curve for testing of the combined set using MiRTM.

The use of transcript ratios has been previously demonstrated to strengthen the prediction accuracy through simplification of features (Avissar et al. 2009; Reddy et al. 2015). We developed a diagnostic score using all ratios of melanoma-enriched miRNAs to melanoma-depleted miRNAs (Fig. 2a). This approach controls for variations in lesion composition (e.g. the relative amount of malignant and benign tissue) when micro-dissection boundaries are not known and differences in tumor cellularity when genetic data are not available. This approach also normalizes for the fraction of miRNAs of melanocytic origin (as opposed to the totality of miRNAs), amplifies the signal from malignant cells by normalizing melanoma-enriched miRNAs to melanoma-depleted miRNAs, and permits cross-platform comparisons without the need for cross-platform normalization. We divided each of the two melanoma-enriched miRNAs by each of the four melanoma-depleted miRNAs, producing eight miRNA ratios (miR-31-5p/ miR-211-5p, miR-31-5p/ miR-125a-5p, miR-31-5p/ miR-125b-5p, miR-31-5p/ miR-100-5p, miR-21-5p/ miR-211-5p, miR-21-5p/ miR-125a-5p, miR-21-5p/ miR-125b-5p, and miR-21-5p/ miR-100-5p). These eight ratios were used to train a random forest classifier. The final miRNA Ratio Trained Model (MiRTM) resulted in an area under receiver operating characteristic curves (AUC) of 1.0 for the discovery set of 7 samples containing 15 matched melanoma and nevus regions.

### Validation of the MiRTM on previously published datasets

To test the accuracy of the MiRTM on independent datasets we obtained and combined the raw data from five previously published miRNA profiling studies (Figs. S1, Table S1; Sand, Xu, Chen, Komina & Jukic). These studies contained both microarray and RT-qPCR datasets. Regardless of the platform, the eight miRNA expression ratios were used as input for the model trained on the sequencing data. The MiRTM resulted in an AUC of ROC of 0.92 (Fig. 2d) and an AUC of Precision-Recall of 0.95 (Fig. 2e). We set two thresholds to calculate sensitivity and specificity. First, we used the optimal threshold as defined by our sequencing cohort (0.5), which resulted in a sensitivity of 0.83 and a specificity of 0.83 (Fig. 2d, red point). As this validation cohort was constructed using different technical platforms for miRNA profiling, we also calculated sensitivity and specificity using the optimal threshold for the ROC curve as 0.94 and 0.83 respectively (Fig. 2d, blue point).

To further examine the reproducibility of our model, we ran MiRTM on each individual previously published dataset. Across all datasets, the MiRTM resulted in an AUC of 0.8 or higher, with an average of 0.922 (Fig. S5a). Optimal sensitivity remained above 0.9 across each dataset (Fig. S5b). With the exception of one dataset, optimal specificity was above 0.8 (Fig. S5b). Interestingly, that dataset (Fig. S5, X) contained the highest level of the miRNA network we identified as associated with tumor cellularity (Fig. 1e, X).)

### Validation of the MiRTM on randomly selected cases

Discovery phase cohorts are often selected for unambiguous and homogenous cases. To further validate our model on a greater diversity of cases, we randomly retrieved 82 biopsied melanocytic lesions - 41 neoplasms diagnosed as nevi and 41 diagnosed as melanoma - from the archives of the UCSF Dermatopathology Section. All diagnoses were reviewed and confirmed by an independent dermatopathologist. This cohort contained a greater range of tumor cellularity and subtypes of melanocytic neoplasms than the discovery cohort (Table 1, Table S5). Instead of micro-dissection, entire FFPE sections were scraped to obtain bulk RNA. The abundance of the six miRNAs was assessed by RT-qPCR (Fig. 3a). We converted C_t_ values to expression ratios using the linear transformation (2^ -C_t_) and ran MiRTM. The resultant AUC of ROC for our unfiltered cohort (UC) was 0.92 (Fig. 3b). The AUC of the Precision-Recall curve was 0.911 (Fig. 3c). To classify the lesions into benign and malignant, we again considered two thresholds. Using the threshold defined by our sequencing cohort (0.5) the model achieved sensitivity of 0.83 and specificity of 0.71 (Fig. 3b red point). The optimal sensitivity and specificity of this ROC curve were 0.81 and 0.90 (Fig. 3b blue point).

**Table 1:**
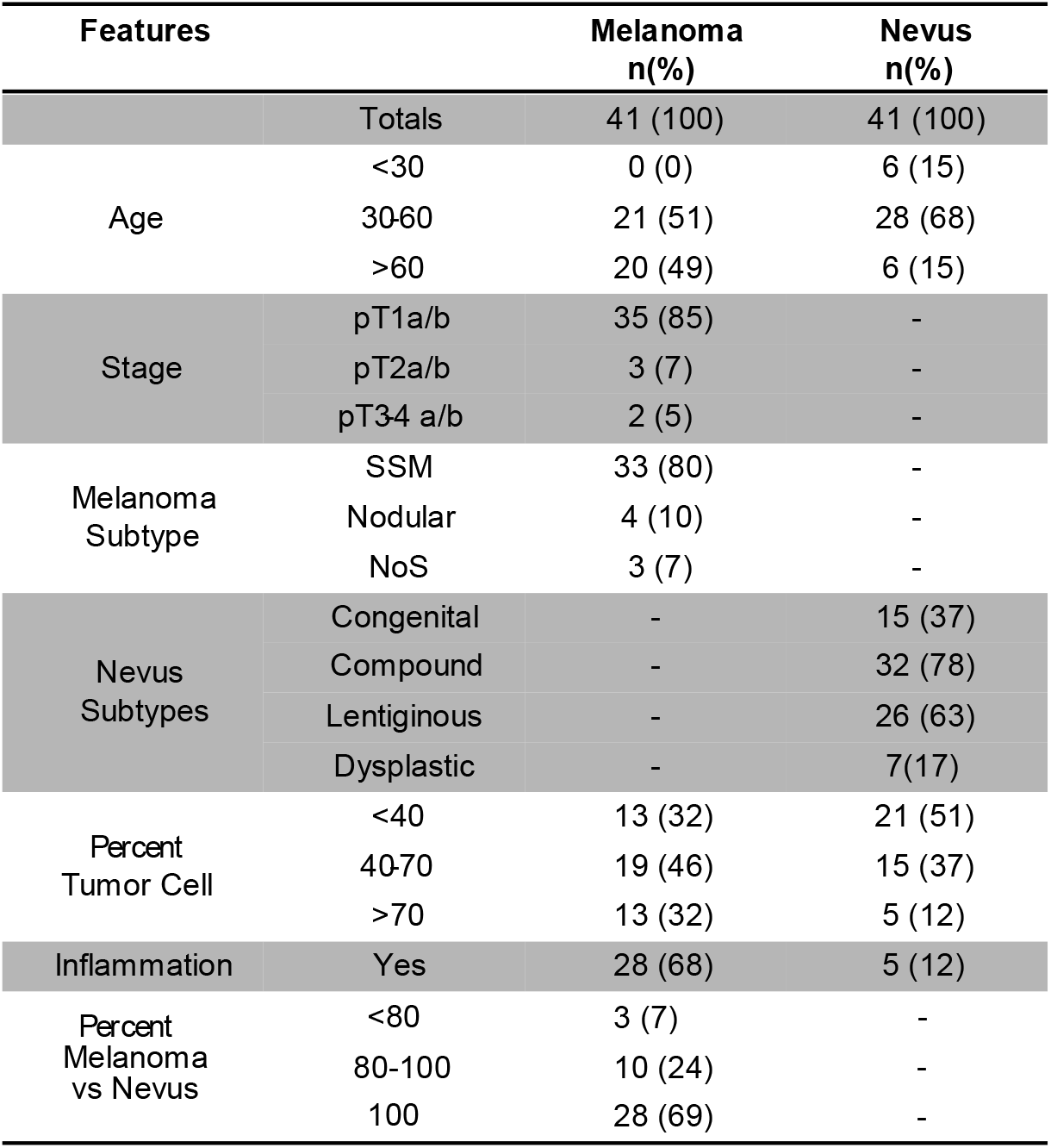
Sample information for unfiltered cohort.

| Features |  | Melanoma<br>n(%) | Nevus<br>n(%) |
| --- | --- | --- | --- |
| Totals |  | 41 (100) | 41 (100) |
| Age | <30 | 0 (0) | 6 (15) |
|  | 30-60 | 21 (51) | 28 (68) |
|  | >60 | 20 (49) | 6 (15) |
| Stage | pT1a/b | 35 (85) | - |
|  | pT2a/b | 3 (7) | - |
|  | pT3-4 a/b | 2 (5) | - |
| Melanoma<br>Subtype | SSM | 33 (80) | - |
|  | Nodular | 4 (10) | - |
|  | NoS | 3 (7) | - |
| Nevus<br>Subtypes | Congenital | - | 15 (37) |
|  | Compound | - | 32 (78) |
|  | Lentiginous | - | 26 (63) |
|  | Dysplastic | - | 7 (17) |
| Percent<br>Tumor Cell | <40 | 13 (32) | 21 (51) |
|  | 40-70 | 19 (46) | 15 (37) |
|  | >70 | 13 (32) | 5 (12) |
| Inflammation | Yes | 28 (68) | 5 (12) |
| Percent<br>Melanoma<br>vs Nevus | <80 | 3 (7) | - |
|  | 80-100 | 10 (24) | - |
|  | 100 | 28 (69) | - |

**Figure 3:**
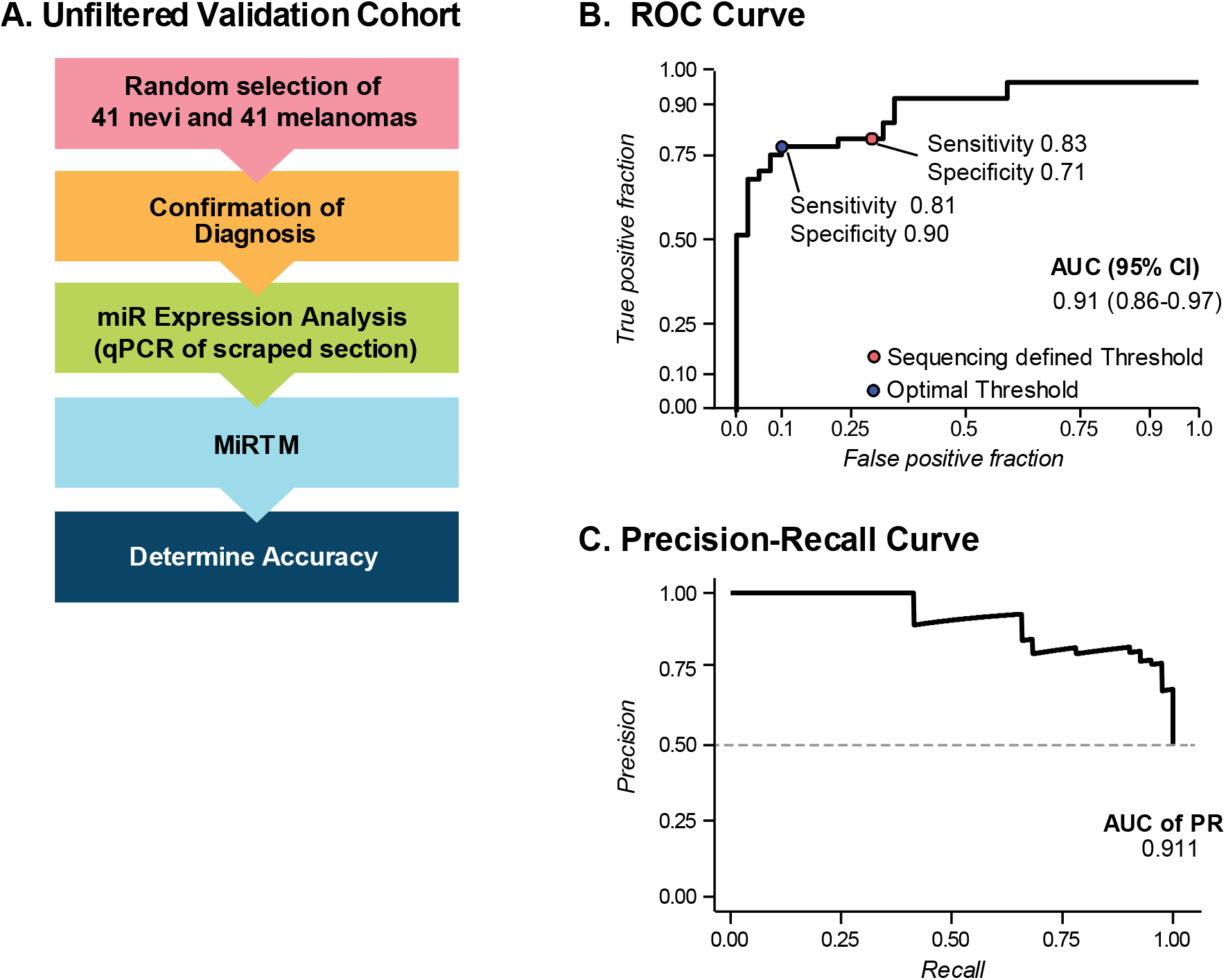
MiRTM validation. A) Workflow for assembling and testing a randomly selected cohort of melanomas and nevi for the validation cohort. B) ROC curve for validation cohort using MiRTM. Threshold determined from the discovery sequencing set and optimal threshold are shown in as red and blue points respectively with corresponding sensitivity and specificity annotated for each. C) Precision-Recall curve for testing of the validation cohort using MiRTM.

As the MiRTM demonstrated a lower sensitivity for the more diverse second validation cohort compared to the previously published cohorts, we examined whether the model was affected by tumor cellularity and patient age. When we compared the MiRTM score to the tumor cellularity in each sample as assessed by a dermatopathologist, we found no correlation (Fig. 4a), suggesting that the MiRTM score was unaffected by this variable. Similarly, we found no correlation between MiRTM score and age (Fig. S6). To investigate whether other features might influence the MiRTM score, we calculated the correlations between fourteen clinical features and the MiRTM score (Fig. 4b-c, Table S5). In the melanoma samples, we observed the expected correlations between clinical features such as age and solar elastosis as a proxy for mutation burden (Fig. 4b). However, we observed no significant correlations with the MiRTM score that suggested any of the features other than diagnosis could influence the score. For example, changes in the thickness or size of the lesion did not affect the MiRTM score. Similarly, in the nevus samples, most features did not influence the score, including the presence of dysplastic features (Fig. 4c). However, exclusively among the benign lesions, the MiRTM score was positively correlated with inflammation (Fig. 4c-d). Together, these data demonstrate that while the MiRTM score is robust against heterogeneity in lesion size, tumor cellularity, and presence of dysplastic features, the presence of inflammatory cells might result in false positive calls.

**Figure 4:**
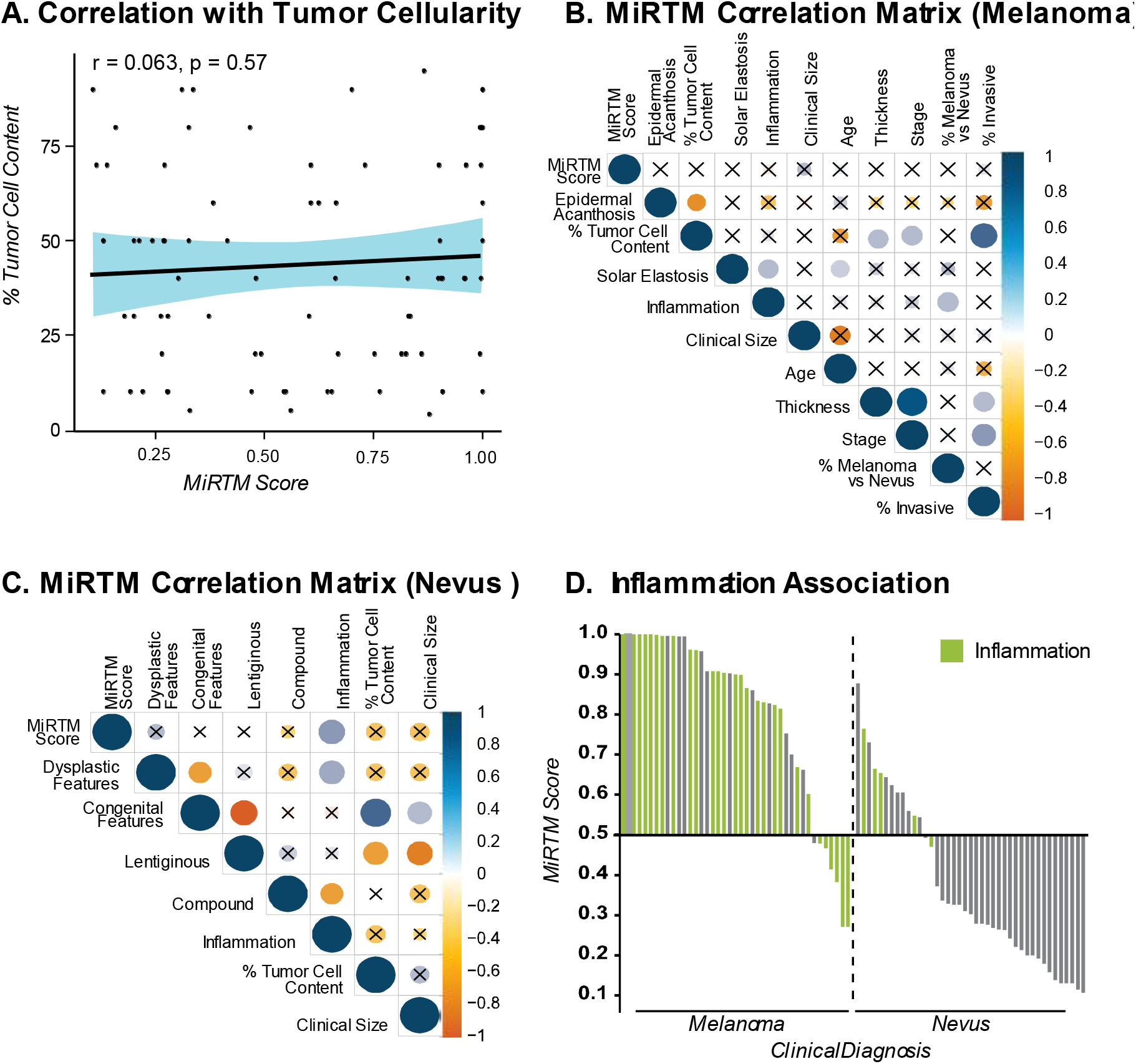
MiRTM accuracy is unaffected by tumor cellularity A) Correlation of MiRTM score with percent tumor cell content. 95% CI is shown in blue. B-C) Correlation matrix for histological and clinical features with MiRTM score for melanoma (B) and nevus (C) samples. Non-significant correlations are shown with an X, while direction and size of significant correlations are indicated by color. D) MiRTM scores for each sample in the validation cohort. Scores above 0.5 were called malignant. Samples microscopically graded as inflamed are colored in green.

## Discussion

Numerous studies have analyzed miRNA expression at different stages of melanoma progression, collectively identifying over 500 miRNAs enriched in nevi or melanomas (Babapoor et al. 2017; Kozubek et al. 2013; Hanniford et al. 2015; Jukic et al. 2010; Satzger et al. 2012; Sand et al. 2013; Xu et al. 2012; Komina et al. 2016; Chen et al. 2011; Latchana et al. 2017). Our analyses have refined this expansive list to six miRNAs that reproducibly distinguish nevi from melanoma across independent datasets and profiling platforms. We identified this signature by controlling two important variables, interobserver variability of diagnosis and variability in tumor cellularity. To address the first variable, we utilized a discovery cohort for which the diagnosis accuracy was established by requiring the median concordance among five to eight different dermatopathologists and supporting genetic features, eliminating the possibility of training the model with misdiagnosed cases. Secondly, melanomas and nevi were matched as pairs representing different progression stages of the same neoplastic clone, eliminating variability by comparing different lesions from different patients. Thirdly, we identified and excluded miRNAs from the signature, whose expression is influenced by tumor cellularity, thereby removing a covariate that has confounded previous analyses. When tested on six datasets assembled by independent groups, a model trained on expression ratios of the refined signature classified benign from malignant melanocytic lesions with an average AUC of ROC above 0.91.

Our strategy required a meticulously assembled and annotated initial cohort of lesions and next generation small RNA sequencing. Although this approach permitted us to identify confounding covariates, the cost and effort to obtain each case also constrained the size of the training cohort to only fifteen samples. One fundamental feature of machine learning is that increased training improves accuracy. Thus, the current performance of the model represents the lower bound of the potential accuracy that could be obtained with a larger training of similar cases. Despite the limited size of the training set, the sensitivity and specificity of the MiRTM for validation sets thresholded on the discovery cohort was 0.83 and 0.71-0.83. This performance of the MiRTM is comparable to other molecular tests for distinguishing benign melanocytic nevi from melanoma, including chromosomal analysis by fluorescence in situ hybridization (sensitivity 0.72-1.00, specificity 0.90-1.00) (Gerami et al. 2010; Ferrara and De Vanna 2016) and myPath Melanoma gene expression profiling (sensitivity 0.63-0.90, specificity 0.88-0.93)(Clarke et al. 2017; Minca et al. 2016). The MiRTM does not perform as well as chromosomal analysis by array comparative genomic hybridization (aCGH, sensitivity 0.92-0.96, specificity 0.87-1.00)(Bastian et al. 2003; Wang et al. 2013). However, assessment by the MiRTM requires only a single section of FFPE material, does not require microdissection and RT-qPCR is a quick and affordable assay making this approach a candidate for lesions where tissue availability is limited.

It is important to note that our discovery set and validation sets were obtained using diverse platforms of miRNA profiling. Indeed, the threshold for optimal thresholds for both validation sets (microarray and RT-qPCR) were slightly different than the optimal threshold for the discovery set (small RNA sequencing). Although the over-all crossplatform performance demonstrates the robustness of the model, future studies aimed at clinical development of this model should consider training and validating the model on a single platform.

The only measured feature that correlated with the MiRTM score overall was diagnosis. However, in the benign samples of the validation set, we observed that inflammation can result in false-positive calls. Indeed, some of the miRNAs (miR-125b, miR-31, miR-21) have been associated with inflammation in psoriasis (Hawkes et al. 2016). However, inflammation was not correlated with the MiRTM score over all samples in the cohort suggesting the feature-selected miRNAs are not exclusively an inflammation signature. Further training on a larger cohort selected for differential inflammation status could substantially reduce the number of false positives.

Of the six miRNAs of our signature, three (miR-211-5p, miR-21-5p, and miR-125b-5p) (Fig. S1a) have been linked to melanoma, have been previously validated by in situ hybridization (Babapoor et al. 2016; Latchana et al. 2016), and have been functionally assessed in melanoma cell lines. MiR-21 is an established oncomir and regulates genes involved in increased proliferation and invasion (Satzger et al. 2012). It is upregulated in many cancers including melanoma and its expression correlates with progression from nevi to primary melanomas and then to metastatic melanomas (Satzger et al. 2012; Jiang et al. 2012). Conversely, miR-125b is often downregulated in cancers, including advanced melanomas, where its loss results in increased expression of cJUN and MLK3 (Zhang et al. 2014; Kappelmann et al. 2013). MiR-211 is among the most well-established functional miRNAs in melanocytes and is downstream of the important melanocyte lineage transcription factor MITF (Mazar et al. 2010). It is often downregulated during melanoma progression and has been linked to invasion through regulation of BRN2, NFAT5 and TGFβR2 (Levy et al. 2010; Boyle et al. 2011). The other miRNAs in the signature are less well characterized in melanocytes. As another miR-125 family member, miR-125a is expected to target a similar set of genes as miR-125b, but has been mostly described in other cancers. MiR-31 is upregulated in some cancers, but its role as an obligate oncomir is controversial as it is transcribed from a commonly deleted or methylated genomic region in many cancers (Valastyan and Weinberg 2010; Asangani et al. 2012). Similarly, miR-100 has also been described as both a tumor suppressor and an oncomir depending on the context (Li et al. 2015). Regardless of their precise functional role in the context of melanocytic neoplasia, our analyses demonstrate that the relative expression ratios of these six miRNAs can assist in distinguishing benign melanocytic nevi from malignant melanoma in FFPE samples.

## Methods

### Meta-analysis

For meta-analyses summarized in Fig. S1, we used all datasets in public databases that contained miRNA profiling for both primary melanoma and nevus samples for comparison to our knowledge (Table S1) (GSE19229, GSE36236, GSE24996, GSE62372, GSE35579, GSE34460, and E-MTAB-4915). The top differentially expressed miRNAs for each dataset were determined using an FDR cutoff of 0.05 using either Limma (microarray and TaqMan array data (Ritchie et al. 2015)) or DeSeq2 (miRNA-seq data (Love et al. 2014)). To determine overlap (Fig. S1a), only those miRNAs for which probes were included in every detection platform were considered (Table S2). Overlap was plotted using the UpsetR package in R (Conway et al. 2017).

### Clinical specimens and histopathologic assessment

A training cohort of melanomas with an intact adjacent benign nevus constituted the discovery cohort for this study. Fifteen different areas (8 malignant and 7 benign) from seven cases were selected based upon which samples from a larger published cohort that was previously genetically assessed (Shain et al. 2018) had leftover material. All cases had previously been retrieved from the UCSF Dermatopathology archive as formalin-fixed paraffin-embedded (FFPE) tissue blocks. Histopathologically distinct areas had been independently evaluated by a panel of 5-8 dermatopathologists for staging (Shain et al. 2018). Distinct tumor areas were manually micro-dissected with a scalpel under a dissection scope from unstained tissue sections following the guidance of a pathologist in order to limit stromal cell contamination. Previously, genetic DNA had been isolated from four 10μM sections using Qiagen DNA FFPE Tissue Kit (Cat# 56404) (Shain et al. 2018). For this study, four additional 20uM sections were dissected and total RNA was isolated using the RecoverALL Total Nucleic Acid Isolation Kit for FFPE (Ambion).

An independent validation cohort was generated by retrieving obtaining 82 diagnosed melanomas (41 cases) or nevi (41 cases) from UCSF Dermatopathology. Cases were reevaluated by a separate dermatopathologist to confirm diagnosis and obtain histopathological features, but were not excluded for any reason. For RNA isolation of the test cohort, one 20 μM section was scraped off the slide and processed in its entirety without micro-dissection and total RNA was isolated using the RecoverALL Total Nucleic Acid Isolation Kit for FFPE (Ambion).

### MicroRNA-seq and analysis

MicroRNA sequencing libraries were constructed with the TailorMix Small RNA Library Preparation Kit (SeqMatic, CA) using total RNA extracted from FFPE samples. Sequencing was performed on the Illumina HiSeq2500 platform at single-end 50bp. After adaptor sequences were removed, reads were aligned to a human reference (hg37) with Bowtie (Langmead et al. 2009) and then small RNA reference groups (miRBase21) were counted. Data were submitted to dbGaP (phs001550.v2.p1). Differential expression analysis was performed from feature counts using DeSeq2 (Love et al. 2014) with p-values adjusted for multiple testing with the Benjamin-Hochberg method (p-adj).

### Co-expression Analysis

Co-expression analysis was restricted to 805 miRNAs with at least one read in at least two samples. Three co-expression networks were identified in R using a four-step approach as previously described (Lui et al. 2014) and a minimum 10-member seed and 0.85 correlation threshold. First, pairwise biweight midcorrelation coefficients (cor) were calculated for all possible pairs of miRNAs for all samples. Second, miRNAs were clustered using the flashClust implementation of a hierarchical clustering procedure with complete linkage and 1 – cor as a distance measure (Langfelder and Horvath 2008). The resulting dendrogram was cut at a static height of ~0.48, corresponding to the top 10% of pairwise correlations for the entire dataset. Third, all clusters consisting of at least 10 members were identified and summarized by their eigengene (i.e. the first principal component obtained via singular value decomposition of the standardized miRNA expression matrix corresponding to each initial cluster) (Horvath and Dong 2008). Fourth, highly similar networks were merged if the Pearson correlation coefficients of their eigengenes exceeded 0.85. This procedure was performed iteratively such that the pair of networks with the highest correlation > 0.85 was merged, followed by recalculation of all eigengenes, followed by recalculation of all correlations, until no pairs of networks exceeded the threshold. Following these steps, three co-expression networks were identified. The strength of association (*k*_ME_) between each miRNA and each network was determined by calculating the Pearson correlation between its expression pattern over all samples with each eigengene (Horvath and Dong 2008) (Table S4).

### Linear Discriminant Analysis

To assess and visualize the degree to which the three different miRNA groups can be distinguished based on their expression in our samples, we constructed a classifier that predicts the miRNA network. We used Linear Discriminant Analysis to project the expression of miRNAs in each sample into a two-dimensional linear subspace that optimally separates the different miRNA categories. We subsequently trained a support vector machine with the linear kernel to distinguish between the miRNA categories. An area under the ROC curve for distinguishing blue, green and orange categories (Fig. 1c) in a testing set was 0.97, 0.94, and 0.95, respectively.

### Gene set enrichment analysis

GSEA was conducted using the three co-expressed miRNA networks as gene sets against public miRNA expression datasets (GSE16368) comparing either primary human melanocytes (GSM817251 GSM1127159, GSM1127164) to keratinocytes (GSM817253, GSM1127111, GSM1127113) or primary human melanocytes to fibroblasts (GSM817252, GSM1127116). Positive enrichment of each case corresponded to melanocyte-enriched and negative corresponded to either fibroblast- or keratinocyte-enriched.

### Genomic Analysis

The targeted exon sequencing datasets for each sample of the training cohort were accessed from dbGaP (phs001550.v1.p1). Read alignment, mutational analysis, and copy number analysis was performed as previously described (Shain et al. 2015; Talevich et al. 2016). Briefly, sequences were aligned using Burrows-Wheeler Aligner (BWA) (Li and Durbin 2009) with mutational analysis and processing performed using Picard and Genome Analysis Toolkit. Copy number information was obtained with the use of CNVkit. Tumor cell content (tumor cellularity) was calculated bioinformatically using multiple methods when possible, including median mutant allele frequency (MAF) of somatic mutations, MAF of the driver mutation, allelic imbalance over germline and others as previously described (Shain et al. 2018).

### Classifier analyses

A feature subset was selected using the Boruta R package (Kursa and Rudnicki 2010) to determine a minimal set of miRNAs for classifier predictive accuracy from the FFPE miRNA-seq data set. Briefly, for each feature (miRNA) present in the RNA-seq data set, a control “shadow-feature” (shadow miR) of comparable expression and variance was generated through random re-assignment of the read counts to different samples (Fig. S3). The combined feature set (miRNAs and shadow miRs) was used to train a random forest classifier and the importance of each feature for the accuracy of the model was determined. This process was repeated in 1000 iterations, with miRNAs excluded from each ensuing round once they were significantly less important than the maximum important shadow miR. Thus, the remaining list of miRNAs at the conclusion of the analyses represents only those miRNAs that out-performed an equal number of randomized controls by a statistically significant margin. This initial miRNA list was then further refined by removing miRNAs that were below a minimum expression threshold and/or were not detected across all outside test sets to obtain a final list of 6 miRNAs (miR-211-5p, miR-125a-5p, 125b-5p, miR-100-5p, miR-31-5p and miR-21-5p). Using log fold-change information from differential expression analysis, each miRNA was associated as melanoma-enriched (ME) or melanoma-depleted / nevus-associated (MD) and miRNA ratios were created from each combination of the 2 ME miRNA (miR-31-5p and miR-21-5p) and 4 MD miRNA (miR-211-5p, miR-125a-5p, 125b-5p, miR-100-5p). A random forest classifier was then built using this transformed minimal signature set and tested by 5-fold repeated cross-validation over 100 repeats to create a final miRNA ratio trained model (MiRTM). The MiRTM was used to classify the datasets described in the meta-analysis where sufficient data were available to obtain sensitivities, specificities, and overall performance by AUC through a ROC curve for each set (Fig. S5) or combined as a group (Fig. 2d-e). The Dadras and Hernando datasets were omitted from the analysis due to insufficient sample sizes (Dadras contained 2 nevus samples), or too many missing features and large sample imbalance (Hernando contained a nevus/mel ratio of 0.1 and many features removed in processing) (Kozubek et al. 2013; Hanniford et al. 2015). Similarly, MiRTM was used to classify each case in our validation set with sensitivities and specificities determined using either an optimal threshold based on the Youden index or the sequencing determined threshold 0.5 (Fig. 3b). Overall performance was visualized by the area under a ROC or precision recall curve respectively (Fig. 3b-c).

### miRNA qPCR assay

Total RNA was converted into cDNA using the TaqMan Advanced miRNA cDNA Synthesis Kit (Thermo Fisher A28007) following the manufacturer recommended protocols. Quantitative PCR for specific miRNA detection was conducted with TaqMan Advanced miRNA Assays (Thermo Fisher) using the TaqMan Fast Advanced Master Mix (Thermo Fisher 4444557) and analyzed on the Applied Biosystems 7900HT instrument following recommended protocols.

### Statistical Analysis

Statistical significance was set to 0.05 with p-values adjusted for multiple testing with the Benjamin-Hochberg method. Pearson correlation coefficients were obtained between all continuous features with the equivalent point biserial correlation coefficient for binary variables. Correlation matrices were plotted with the corrplot R Package and correlation plots with the ggpubr R package with 95% confidence intervals calculated for the curves. Sensitivities and specificities were calculated from classification models built using the caret R package. ROC curves were generated using the pROC R package. Confidence intervals (CI) were calculated from the 95% CI of 2000 bootstrap replicates for sensitivity and specificity or the ‘Delong’ method for AUCs using pROC R package. The precision-recall plot was generated using the precrec R package. All data was processed in R (3.3.2)

## Data Access

The sequrncing data was submitted to the NCBI database of Genotypes and Phenotypes (dbGaP) accession number phs001550.v2.p1.

## Acknowledgements

This research was supported in part by NIH DP5OD019787 to R.L.J., the Sandler Foundation Program for Breakthrough Biomedical Research Fellowship to R.L.J., the Marcus Program in Precision Medicine Innovation Fund Seeding Big Ideas Award to R.L.J and M.L.W. and the Helen Diller Family Comprehensive Cancer Center Impact Award to M.L.W. and R.L.J. We would like to thank Tim H. McCalmont for assistance in identifying cases and Hilary Faith Hickman for assistance in visual design.

## Author Contributions

R.T. conducted meta-analyses, gene set enrichment analyses, feature selection and training of classifiers. M.H. performed the LDA. S.J.S and M.C.O. performed the co-expression network analyses. Cohort 1 was assembled by A.H.S. I.Y., B.C.B. and N.M.J and micro-dissected by N.M.J and R.L.J. R.L.J. conducted RNA and DNA isolation. A.H.S. performed genetic analyses. Cohort 2 samples and features were assembled by U.E.L. R.T. conducted RNA isolation, RT-qPCR, and analyses. U.E.L., N.M.J, I.Y., and B.C.B. provided histopathology and diagnoses. R.T., I.Y., M.L.W., M.C.O., B.C.B and R.L.J provided significant intellectual contribution and critical reading of the manuscript. R.L.J. conceived of and directed all experiments.

